# Combined Ventilation Of Two Subjects With A Single Mechanical Ventilator Using A New Medical Device: An *In Vitro* Study

**DOI:** 10.1101/2021.01.06.425652

**Authors:** Ignacio Lugones, Matías Ramos, Fernanda Biancolini, Roberto Orofino Giambastiani

## Abstract

**INTRODUCTION:** The SARS-CoV2 pandemic has created a sudden lack of ventilators. DuplicAR^®^ is a novel device that allows simultaneous and independent ventilation of two subjects with a single ventilator. The aims of this study are: a) to determine the efficacy of DuplicAR^®^ to independently regulate the peak and positive-end expiratory pressures in each subject, both under pressure-controlled ventilation and volume-controlled ventilation, and b) to determine the ventilation mode in which DuplicAR^®^ presents the best performance and safety.

**MATERIALS AND METHODS:** Two test lungs are connected to a single ventilator using DuplicAR^®^. Three experimental stages are established: 1) two identical subjects, 2) two subjects with the same weight but different lung compliance, and 3) two subjects with different weight and lung compliance. In each stage, the test lungs are ventilated in two ventilation modes. The positive-end expiratory pressure requirements are increased successively in one of the subjects. The goal is to achieve a tidal volume of 7 ml/kg for each subject in all different stages through manipulation of the ventilator and the DuplicAR^®^ controllers.

**RESULTS:** DuplicAR^®^ allows adequate ventilation of two subjects with different weight and/or lung compliance and/or PEEP requirements. This is achieved by adjusting the total tidal volume for both subjects (in volume-controlled ventilation) or the highest peak pressure needed (in pressure-controlled ventilation) along with the basal positive-end expiratory pressure on the ventilator, and simultaneously manipulating the DuplicAR^®^ controllers to decrease the tidal volume or the peak pressure in the subject that needs less and/or to increase the positive-end expiratory pressure in the subject that needs more. While ventilatory goals can be achieved in any of the ventilation modes, DuplicAR^®^ performs better in pressure-controlled ventilation, as changes experienced in the variables of one subject do not modify the other one.

**CONCLUSIONS:** DuplicAR^®^ is an effective tool to manage the peak inspiratory pressure and the positive-end expiratory pressure independently in two subjects connected to a single ventilator. The driving pressure can be adjusted to meet the requirements of subjects with different weight and lung compliance. Pressure-controlled ventilation has advantages over volume-controlled ventilation and is therefore the recommended ventilation mode.

## INTRODUCTION

As a result of major disasters, health systems have focused on the need for simultaneous medical care for a large number of victims (1, 2, 3, 4, 5). The global crisis triggered by the SARS-CoV2 pandemic created a sudden shortage of ventilators that has led to an increase in the number of deaths (6). Triage protocols are needed for the efficient allocation of these scarce critical care resources (7,8), considering that this could result in potential legal liability (9).

Since the beginning of the pandemic, several teams have focused on multiple ventilation as a possible solution to this huge problem. However, these efforts were soon discouraged by some of the most important scientific societies due to the limitations and potential risks of this strategy (10).

Recently, we reported the results of the evaluation in lung-healthy pigs of a new medical device called DuplicAR^®^ (11). This device was designed to enable mechanical ventilation of two subjects with a single ventilator, without cross-contamination, and allowing for independent management of the peak inspiratory pressure (PIP) and the positive-end expiratory pressure (PEEP).

The aims of this study are: a) to determine the efficacy of the DuplicAR^®^ device to independently regulate the PIP and PEEP in each subject, both under pressure-controlled ventilation (PCV) and volume-controlled ventilation (VCV), and b) to determine the ventilation mode in which DuplicAR^®^ presents the best performance and safety.

## MATERIALS AND METHODS

### The DuplicAR^®^ device

DuplicAR^®^ is a novel device that allows simultaneous and independent ventilation of two subjects with only one ventilator. It is a simple adapter that is connected to the ventilator and provides independent pressurization of the system for the two subjects (Figure 1). It consists of two adapters embedded in the same device. The inspiratory part of the device connects the inspiratory port of the ventilator to each subject’s inspiratory limb and has: 1) valves that regulate flow, and therefore tidal volume (VT) and PIP, and 2) one-way valves that allow independent management of the two circuits and avoid cross-contamination. The expiratory part of the device connects to each subject's expiratory limb and to the expiratory port of the ventilator. Each expiratory limb has a PEEP controller for independent management of this variable. Cross-contamination is prevented through one-way valves and microbiological filters in each line of the circuit.

**Figure 1:**
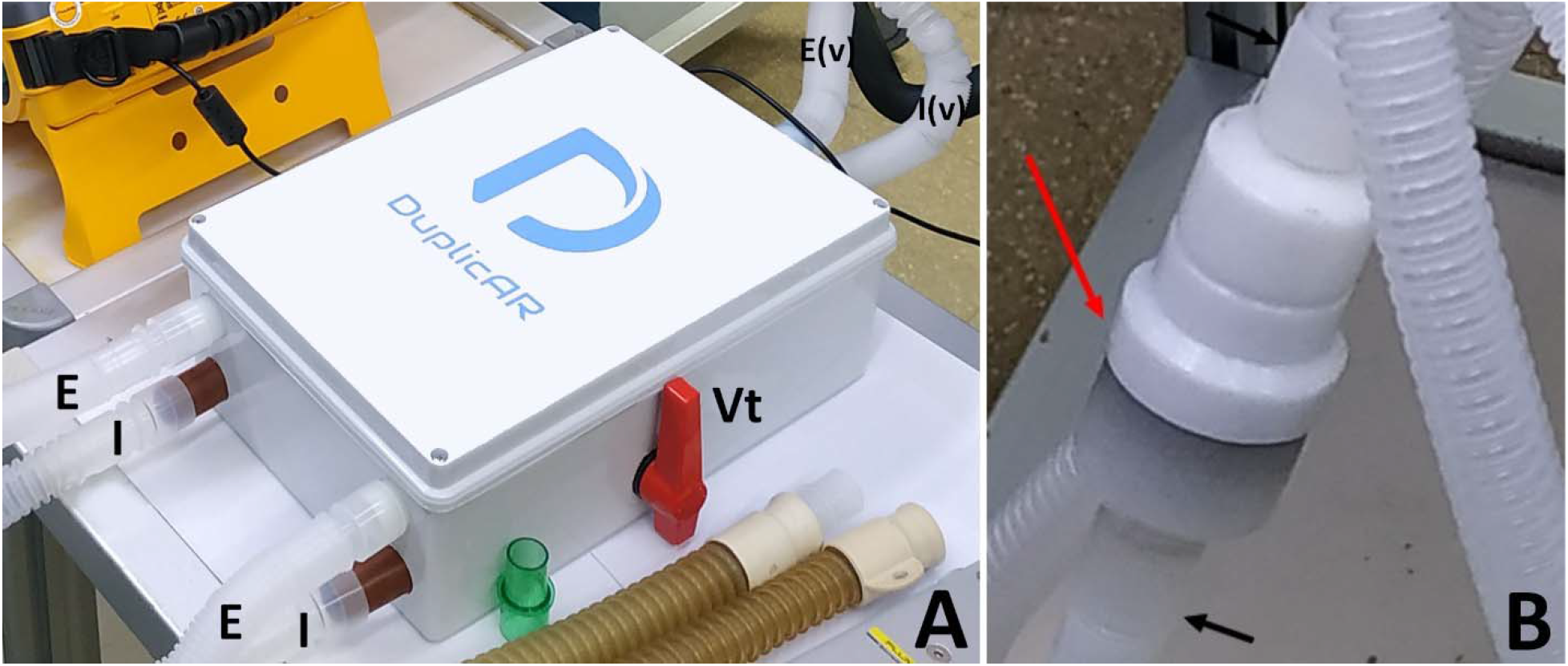
The DuplicAR^®^ device: A) the device with its connections; B) the PEEP valve controller (E: expiratory limb of each subject; E(v): expiratory limb connected to the ventilator; I: inspiratory limb of each subject; I(v): inspiratory limb connected to the ventilator; Vt: tidal volume/peak-pressure controller; red arrow: PEEP controller; black arrows: expiratory limb).

### Experimental model

The effectiveness of the DuplicAR^®^ device for combined ventilation is studied in an *in vitro* experimental model (Figure 2). Two artificial lungs (test lungs), from now on called Subject A and Subject B, are connected to a single ventilator using the DuplicAR^®^ device. Subject A is considered “standard”. Its attributes remain unchanged throughout the experiments. It simulates a 50 kg patient and is represented by a test lung with a compliance of 30 ml/cmH_2_O (0.6 ml cmH_2_O^−1^ kg^−1^). The target VT is 350 ml (7 ml/kg). On the other hand, Subject B is modified throughout the experiments to represent different scenarios. A VT normalized to weight of 7 ml/kg is also targeted for this subject.

**Figure 2:**
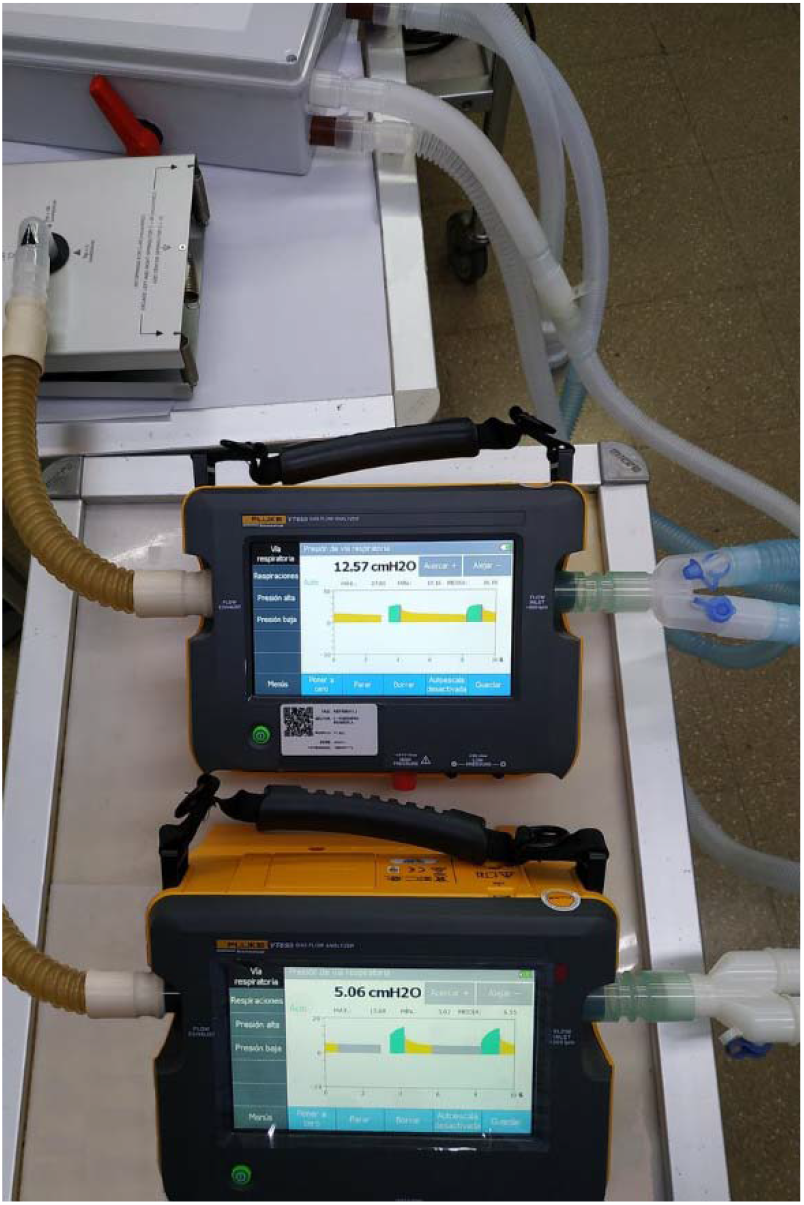
The experimental model: a ventilator connected through the DuplicAR^®^ device to two gas flow analyzers and their respective test lungs.

In all cases, the respiratory rate is a fixed parameter (12 ventilations per minute), which cannot be individualized for each subject. We assume that ventilation occurs in the region between the lower and upper inflection points of the compliance curve. According to this, the compliance of each subject is modeled as a constant. Both subjects are presumed to be conditioned for their best lung compliance curve. In clinical terms, this would mean that patients’ lungs have been optimally recruited.

The experiments are performed using a Newport e360 ventilator (Newport Medical, Costa Mesa, CA) connected to the DuplicAR^®^ system. This device is in turn connected to two portable precision test lungs with adjustable compliance and resistance (ACCU LUNG, Fluke Biomedical, Everett, WA, USA) using regular circuit tubing. Two gas flow analyzers (VT 650, Flukke Biomedical, Everett, WA, USA) are also used, each one placed between the test lung and its respective Y-piece (Figure 2).

### Experimental stages

Three experimental stages are established for the study, according to the simulated patient’s weights and lung compliances:

- Stage 1: Two identical subjects (the same weight and compliance).
- Stage 2: Two subjects with the same weight but different compliance.
- Stage 3: Two subjects with different weight and compliance.

At each stage, subjects A and B are connected to the ventilator using the DuplicAR^^®^^ device and ventilated in PCV and VCV. The target volume established for each subject in all stages is 7 ml/kg. This target VT is achieved through manipulation of the ventilator (PIP in PCV or VT in VCV) in combination with the flow controllers of the DuplicAR^®^ device. The PEEP in the ventilator is set to 5 cmH_2_O and maintained at that value throughout the experience. In *Experiment I*, considered the “baseline situation”, there is no difference in PEEP among subjects. A difference in PEEP is achieved through manipulation of the PEEP controller of the DuplicAR^®^ device. The PEEP in Subject B is then successively increased to 7-10 (*Experiment II*) and 11-14 cmH_2_O (*Experiment III*), while Subject A maintains in 5 cmH_2_O.

The three stages are summarized in Table 1.

**Table 1:** Summary of the three stages, simulating the different possible combinations of subjects’ weights and lung compliances. The ventilator is configured in each mode to adequately ventilate the two subject with a target volume of 7 ml/kg. The controllers of the DuplicAR^®^ device are manipulated to deliver asymmetric flow or different PEEP to the subjects.

|  | Stage 1 |  | Stage 2 |  | Stage 3 |  |
| --- | --- | --- | --- | --- | --- | --- |
|  | A | B | A | B | A | B |
| Weight (kg) | 50 | 50 | 50 | 50 | 50 | 28 |
| Compliance (ml/cmH<sub>2</sub>O) | 30 |  | 30 | 20 | 30 | 10 |
| Compliance (ml cmH<sub>2</sub>O-1 kg-1) | 0.6 |  | 0.6 | 0.4 | 0.6 | 0.35 |
| PEEP ventilator (cmH<sub>2</sub>O) | 5 |  | 5 |  | 5 |  |
| I:E ratio | 1:2 |  | 1:2 |  | 1:2 |  |
| Target volume (ml/kg) | 7 |  | 7 |  | 7 |  |
| Total target load (ml) | 350 | 350 | 350 | 350 | 350 | 196 |
| Peak pressure in PCV (cmH<sub>2</sub>O) | 15 |  | 21 |  | 22 |  |
| Tidal volume in VCV (ml) | 700 |  | 700 |  | 560 |  |
| DuplicAR flow manipulation | No | No | Yes | No | Yes | No |
| PEEP Experiment I (cmH<sub>2</sub>O) | 5 |  | 5 |  | 5 |  |
| PEEP Experiment II (cmH<sub>2</sub>O) | 5 | 10 | 5 | 10 | 5 | 10 |
| PEEP Experiment III (cmH<sub>2</sub>O) | 5 | 15 | 5 | 15 | 5 | 15 |

## RESULTS

### Stage 1

The data are shown in Tables 2 and 3.

**Table 2:** Stage 1 in PCV.

|  | Experiment I |  | Experiment II |  | Experiment III |  |
| --- | --- | --- | --- | --- | --- | --- |
|  | A | B | A | B | A | B |
| PA max (cmH<sub>2</sub>O) | 16.12 | 15.92 | 16.25 | 21.78 | 16.12 | 24.61 |
| PA min (cmH<sub>2</sub>O) | 5.37 | 5.11 | 4.92 | 10.35 | 4.24 | 13.41 |
| Driving (cmH<sub>2</sub>O) | 10.75 | 10.81 | 11.33 | 11.43 | 11.88 | 11.2 |
| C (ml/cmH<sub>2</sub>O) | 33.95 | 33.21 | 31.16 | 30.53 | 30.30 | 30.89 |
| V<sub>t</sub> (ml) | 365 | 359 | 353 | 349 | 360 | 346 |
| V<sub>t</sub>/kg (ml/kg) | 7.3 | 7.18 | 7.06 | 6.98 | 7.2 | 6.92 |

**Table 3:** Stage 1 in VCV.

|  | Experiment I |  | Experiment II |  | Experiment III |  |
| --- | --- | --- | --- | --- | --- | --- |
|  | A | B | A | B | A | B |
| PA max (cmH<sub>2</sub>O) | 15.19 | 15.72 | 15.64 | 22.12 | 15.62 | 24.83 |
| PA min (cmH<sub>2</sub>O) | 5.36 | 5.13 | 5.33 | 10.79 | 4.52 | 13.26 |
| Driving (cmH<sub>2</sub>O) | 9.83 | 10.59 | 10.31 | 11.33 | 11.1 | 11.57 |
| C (ml/cmH<sub>2</sub>O) | 35.10 | 32.86 | 33.56 | 30.45 | 30.27 | 30.51 |
| V<sub>t</sub> (ml) | 345 | 348 | 346 | 345 | 336 | 353 |
| V<sub>t</sub>/kg (ml/kg) | 6.9 | 6.96 | 6.92 | 6.9 | 6.72 | 7.06 |

As evidenced in Experiment I, regardless of the ventilation mode, without intervention, the driving pressure is the same in both subjects (10.5 ± 0.45 cmH_2_0), and therefore, the VT is also the same (354 ± 9.36 ml).

In both ventilation modes, it is possible to increase the PEEP of Subject B without compromising its own driving pressure or that of Subject A. This is achieved in PCV by: 1) increasing the PIP in the ventilator in the same amount as the increase introduced in the PEEP of Subject B and 2) restricting at the same time the PIP of Subject A through manipulation of the flow controller of the DuplicAR^®^ device.

In VCV, as the PEEP of Subject B increases through manipulation of the PEEP controller of the device, it is necessary to adjust the flow controller of Subject A. This is mandatory as otherwise air would not enter Subject B until the system reaches the breakdown pressure generated by the PEEP valve, causing a disproportionate increase in the VT delivered to Subject A.

### Stage 2

The data are shown in Tables 4 and 5.

**Table 4:** Stage 2 in PCV.

|  | Experiment I |  | Experiment II |  | Experiment III |  |
| --- | --- | --- | --- | --- | --- | --- |
|  | A | B | A | B | A | B |
| PA max (cmH<sub>2</sub>O) | 15.78 | 21.57 | 16.01 | 24.72 | 16 | 27.77 |
| PA min (cmH<sub>2</sub>O) | 5.49 | 5.25 | 5.23 | 8.7 | 5.02 | 12.6 |
| Driving (cmH<sub>2</sub>O) | 10.29 | 16.32 | 10.78 | 16.02 | 10.98 | 15.17 |
| C (ml/cmH<sub>2</sub>O) | 34.69 | 21.32 | 33.86 | 22.41 | 31.97 | 23.47 |
| Vt (ml) | 357 | 348 | 365 | 359 | 351 | 356 |
| Vt/kg (ml/kg) | 7.14 | 6.96 | 7.3 | 7.18 | 7.02 | 7.12 |

**Table 5:** Stage 2 in VCV.

|  | Experiment I |  | Experiment II |  | Experiment III |  |
| --- | --- | --- | --- | --- | --- | --- |
|  | A | B | A | B | A | B |
| PA max (cmH2O) | 16.62 | 20.71 | 16.19 | 23.96 | 16 | 27.71 |
| PA min (cmH2O) | 5.46 | 5.22 | 5.11 | 8.54 | 5 | 12.33 |
| Driving (cmH2O) | 11.16 | 15.49 | 11.08 | 15.42 | 11 | 15.38 |
| C (ml/cmH2O) | 31.45 | 21.82 | 30.51 | 22.31 | 30.91 | 21.98 |
| Vt (ml) | 351 | 338 | 338 | 344 | 340 | 338 |
| Vt/kg (ml/kg) | 7.02 | 6.76 | 6.76 | 6.88 | 6.8 | 6.76 |

Both subjects have the same size and require the same VT (350 ml). Since subject B has a decreased compliance compared to Subject A, its driving pressure needs to be higher:

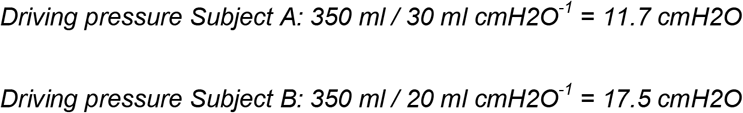

To achieve this difference in each subject’s driving pressure in PCV, the PIP must be increased in the ventilator to deliver the adequate VT to the subject with the lowest compliance (in this case, Subject B) and afterwards, the PIP in Subject A is decreased through manipulation of the respective flow controller of the DuplicAR^®^ device.

In VCV, the VT is delivered by the ventilator to the two subjects as a whole, but its distribution is asymmetric due to the differences in subjects’ characteristics. The adequate VT for each subject needs to be regulated through manipulation of the flow controller of the DuplicAR^®^ device of the subject with greater compliance. In this way, with a single ventilator configuration, different driving pressures are established.

In both ventilation modes, it is possible to modify the PEEP of a subject without compromising its driving pressure or modifying the other subject's parameters. In each of the three experiments, the driving pressures are preserved. In PCV, this is accomplished by increasing the PIP in the ventilator setting to maintain the driving pressure of the subject with an increased PEEP (Subject B). Accurate driving pressure in Subject A is modulated by the flow controller of the DuplicAR^®^ device, thus preventing an excessive PIP (Figure 2). In VCV, the total VT is set. When the PEEP of Subject B is selectively increased, the flow controller of Subject A is adjusted to achieve an adequate distribution of the total VT.

### Stage 3

The data are shown in Tables 6 and 7.

**Table 6:** Stage 3 in PCV.

|  | Experiment I |  | Experiment II |  | Experiment III |  |
| --- | --- | --- | --- | --- | --- | --- |
|  | A | B | A | B | A | B |
| PA max (cmH2O) | 16.15 | 23.7 | 16.09 | 28.92 | 16.84 | 34.1 |
| PA min (cmH2O) | 5.44 | 5.16 | 5.03 | 7.99 | 5 | 11.3 |
| Driving (cmH2O) | 10.71 | 18.54 | 11.06 | 20.93 | 11.84 | 22.8 |
| C (ml/cmH2O) | 35.57 | 10.41 | 31.92 | 9.32 | 32.60 | 9.21 |
| Vt (ml) | 381 | 193 | 353 | 195 | 386 | 210 |
| Vt/kg (ml/kg) | 7.62 | 6.89 | 7.06 | 6.96 | 7.72 | 7.5 |

**Table 7:** Stage 3 in VCV.

|  | Experiment I |  | Experiment II |  | Experiment III |  |
| --- | --- | --- | --- | --- | --- | --- |
|  | A | B | A | B | A | B |
| PA max (cmH2O) | 16.33 | 23.26 | 17.02 | 27.03 | 16.95 | 33.61 |
| PA min (cmH2O) | 5.42 | 5.16 | 5.15 | 7.51 | 4.98 | 11.46 |
| Driving (cmH2O) | 10.91 | 18.1 | 11.87 | 19.52 | 11.97 | 22.15 |
| C (ml/cmH2O) | 30.25 | 10.77 | 30.75 | 9.89 | 29.99 | 9.48 |
| Vt (ml) | 330 | 195 | 365 | 193 | 359 | 210 |
| Vt/kg (ml/kg) | 6.6 | 6.96 | 7.3 | 6.89 | 7.18 | 7.5 |

In this situation, the two subjects have important disparities in terms of weight and lung compliance. Subject A is a simulated patient weighing 50 kg with a lung compliance of 30 ml / cmH2O, while Subject B simulates a patient weighing 28 kg with a lung compliance of 10 ml / cmH2O. The driving pressures that each subject requires are:

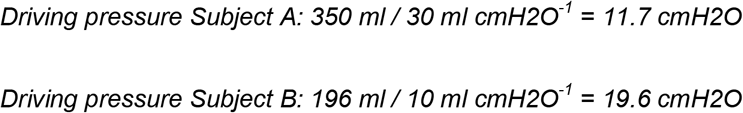

In order to achieve this difference in each subject’s driving pressure in PCV, the PIP in the ventilator must first be configured to ensure ventilation to the subject with the lowest compliance (in this case, Subject B, requiring a PIP of 24 cmH2O). Afterwards, the PIP in Subject A is decreased through manipulation of the respective flow controller of the DuplicAR^®^ device.

In VCV, the different driving pressures of the two subjects are achieved by setting a total VT in the ventilator and managing its distribution among subjects with the flow controllers of the DuplicAR^®^ device, as in Stage 2.

The PEEP is also managed independently for each subject in both ventilation modes. As in Stage 2, in PCV, the driving pressures are preserved by increasing the PIP of the ventilator to maintain the driving pressure of the subject with an increased PEEP (Subject B) and decreasing the PIP of the other subject through manipulation of the respective flow controller of the DuplicAR^®^ device. In VCV, the total VT configured in the ventilator is unequally distributed to meet each subject’s requirements. Since PEEP of Subject B increases, the flow controller of Subject A needs to be adjusted to achieve an adequate VT.

## DISCUSSION

Since the first cases of covid-19 were reported in December 2019 in Wuhan as a cluster of patients with pneumonia, the disease rapidly disseminated worldwide. In a few months, some of the more robust health care systems in the world collapsed, as it was impossible to meet the needs of so many critically ill patients simultaneously. By the end of 2020, more than 80 million people had suffered from the disease and almost 2 million had died.

From the beginning of the pandemic, the shortage of mechanical ventilators was at the center of the scene. People died because resource constraints limited the availability of ventilatory support (6,12). On March 24, 2020, Drs Jeremy Beitler, Richard Branson, and colleagues from the Columbia University College of Physicians and Surgeons and the New York-Presbyterian Hospital published a ventilator sharing protocol for dual-patient ventilation with a single mechanical ventilator for use during critical ventilator shortage (13). This was just one of the multiple responses from the worldwide health care community to the crisis.

However, two days later, on March 26, with 462,680 cases diagnosed worldwide, the Society of Critical Care Medicine (SCCM), the American Association for Respiratory Care (AARC), the American Society of Anesthesiologists (ASA), the Anesthesia Patient Safety Foundation (APSF), the American Association of Critical-Care Nurses (AACN), and the American College of Chest Physicians (CHEST) issued a Joint Statement on the concept of placing multiple patients on a single mechanical ventilator (10). The central message was that sharing mechanical ventilators should not be attempted because it could not be done safely with the equipment available at that time. The document was concise but robust, and addressed all issues related to multiple ventilation as it was being performed at the moment.

At the same time, with the aim of overcoming these issues, our team created a new medical device (patent application submitted), called DuplicAR^®^. This device allows for independent ventilation of two subjects with only one ventilator. In this way, the concept of “combined ventilation” (different from “multiple” or “shared” ventilation) was born. In it, mutual interactions between the two subjects connected to the same ventilator are considered, and individual needs are attended to ventilate them adequately. The results of the use of this device in a lung-healthy animal model were published recently (11).

In the present work, we demonstrate in an *in vitro* model that most of the problems outlined by the Joint Statement can be overcome with this new medical device.

One of the strongest arguments exposed against multiple ventilation was that the VT (or the PIP, according to the mode of mechanical ventilation) and the PEEP (which is of critical importance in covid-19 patients) would be impossible to manage. Here, we demonstrate that these variables can be managed independently to ventilate the two subjects adequately, even if they have different weight and/or lung compliance and/or PEEP requirements. This is achieved by adjusting the total VT for both subjects (or the highest PIP needed) and the basal PEEP on the ventilator and manipulating the DuplicAR^®^ controllers to decrease the VT or PIP in the subject that needs less and increase the PEEP in the subject that needs more.

While we show that the ventilatory goals can be achieved both in PCV and VCV, VCV has major disadvantages compared to PCV. Although the total VT for both subjects in VCV can be distributed asymmetrically between them (using DuplicAR^®^ to overcome the problem of volumes going to the most compliant lung), this mode introduces the possibility of deleterious interactions between subjects. For example, if a subject’s endotracheal tube gets kinked, the other subject will receive a dangerously large VT. This happens because sudden changes in resistance or compliance in one subject directly impact on the other. A similar phenomenon is observed when increasing the PEEP selectively in one subject. In VCV, the other subject receives a greater VT, since it is necessary that the inspiratory tubing reaches greater pressure to allow the volume to be delivered to the subject with greater PEEP. This is accomplished at the expense of increasing volume on the subject with no modification of the PEEP.

All these problems can be overcome using continuous mandatory PCV, the mode of ventilation recommended for DuplicAR^®^, and adding profound sedation and paralyzation of both patients. Despite the fact that a guaranteed VT cannot be delivered to any patient (not different from having only one patient on PCV), the ability to control and limit the driving pressure of each individual regardless of weight, airway resistance and lung compliance may allow this strategy to be reasonably lung-protective. Besides, an increase in one subject's PEEP (without modifying the PIP on the ventilator) only affects the subject in whom the PEEP is modified, by decreasing the driving pressure and therefore the VT. Another advantage of PCV is that sudden changes in one subject (i.e., dynamic compliance variability or endotracheal tube obstruction) do not affect the pressurization of the system, and therefore have no consequences on the other subject (13). Deleterious interactions between patients are then mostly avoided using this mode. For example, if Subject A’s endotracheal tube gets kinked, Subject A will receive a reduced VT, but this will have no impact on Subject B. Finally, PCV spontaneously compensates for the compliance added to the system by the tubing, which in combined ventilation is expected to contain twice the volume compared to single (conventional) ventilation. In VCV, in contrast, this must be manually compensated by adding an extra volume to the total VT delivered by the ventilator.

The individualized management described above can easily be maintained with DuplicAR^®^ over time, as the PIP and PEEP can be adjusted to the subjects’ requirements at any time. Patients could deteriorate and recover at different rates, and the distribution of gas to each patient would be monitored and adjusted to each patient’s needs in a continuous fashion.

Finally, there are two other major concerns highlighted by the Joint Statement, which DuplicAR^®^ can adequately address. The first is related to monitoring and setting alarms during multiple ventilation. In the recently published animal model, we monitor the airway pressures in each animal using a regular pressure transducer connected between the Y-piece and a multiparameter monitor (11). We are currently working on an advanced prototype with alarms, direct measurement in real time of all the variables of each subject and electronic closed-loop control of the PIP and PEEP in each individual through a human-machine interface. This would avoid additional external monitoring.

The second concern outlined by the Joint Statement was that patients could share gas between circuits. Pendelluft was considered possible between patients, leading to both cross-infection and overdistension. However, this is avoided with the DuplicAR^®^ device, which has filters and one-way valves that isolate patient circuits from each other.

## Limitations

The present study has limitations. The test lungs used in this *in vitro* study have constant compliance. In clinical practice, this would imply that regardless of the PEEP and the PIP reached, the relationship between a volume differential and a pressure differential would behave as a constant. In real mechanical ventilation scenarios (and particularly in patients with lung disease), this relationship between volume and pressure may not be linear. In these cases, the VT would be delivered at pressure values close to the “upper inflection point”, and therefore lung compliance would progressively decrease.

In addition, the objective of this experimental *in vitro* model is to represent, in a reductionist way, the interactions between variables in a combined ventilation scenario. This model does not take into account other aspects that should be considered in clinical practice, such as the hemodynamic status, the quality of gas exchange, and the possibility of inadvertent spontaneous ventilation in patients under shared ventilation.

Finally, it is important to note that this device has not yet been tested in humans and its use should only be considered as a life-saving bridge alternative to palliate the consequences of the sudden shortage of ventilators during catastrophic events.

## CONCLUSIONS

DuplicAR^®^ is an effective tool to manage the PIP and PEEP independently in two subjects connected to a single ventilator. The driving pressures can be adjusted to meet the requirements of the two subjects, even if they have different weight, lung compliance, and/or PEEP requirements. DuplicAR^®^ can achieve the ventilatory goals both in PCV and VCV. However, it performs better in continuous mandatory PCV, as the changes experienced in the variables of one subject do not modify the other one. PCV also compensates the compliance changes generated by the additional tubing required in combined ventilation. This new medical device may then become a useful tool to face the shortage of mechanical ventilators during health crises.

## ACKNOWLEDGMENTS

Ignacio Lugones would like to thank Luciano Gentile and Patricia Crego from the Favaloro Foundation University Hospital, Argentina, for supplying equipment to carry out the experiments. This author also thanks Juan José Díaz and Nicolás Campagne from ETYC S.A. for providing technical assistance during the study.

